# Synthetic mucus biomaterials synergize with antibiofilm agents to combat *Pseudomonas aeruginosa* biofilms

**DOI:** 10.1101/2024.08.09.607383

**Authors:** Sydney Yang, Alexa Stern, Gregg Duncan

## Abstract

Bacterial biofilms are often highly resistant to antimicrobials causing persistent infections which when not effectively managed can significantly worsen clinical outcomes. As such, alternatives to standard antibiotic therapies have been highly sought after to address difficult-to-treat biofilm-associated infections. We hypothesized a biomaterial-based approach using the innate functions of mucins to modulate bacterial surface attachment and virulence could provide a new therapeutic strategy against biofilms. Based on our testing in *Pseudomonas aeruginosa* biofilms, we found synthetic mucus biomaterials can inhibit biofilm formation and significantly reduce the thickness of mature biofilms. In addition, we evaluated if synthetic mucus biomaterials could work synergistically with DNase and/or α-amylase for enhanced biofilm dispersal. Combination treatment with these antibiofilm agents and synthetic mucus biomaterials resulted in up to 3 log reductions in viability of mature *P. aeruginosa* biofilms. Overall, this work provides a new bio-inspired, combinatorial approach to address biofilms and antibiotic-resistant bacterial infections.

## Introduction

Chronic infections are often linked to biofilms which form when bacteria attach to surfaces and form a matrix^1^. Opportunistic pathogens such as *Pseudomonas aeruginosa* are often found at chronic wound sites due to this propensity to form biofilms ^2,3^. Biofilm associated infections have high resistance to antibiotic treatments due to the composition of the biofilm matrix which is comprised of extracellular DNA and extracellular polymeric substances (EPS), including bacterial polysaccharides and proteins, that encapsulate bacteria^4–6^. This biofilm matrix forms a barrier protecting bacteria within the biofilm and increases resistance to antibiotics and the immune system^1^. As a result, biofilms and chronic infections are exceptionally difficult to treat. As a result, therapeutics that can overcome these barriers and retain their efficacy against biofilms are highly sought after. For example, nanomaterials have been developed which can readily penetrate the biofilm matrix to enable delivery of antimicrobials^7–10^. Hydrogel-based biomaterials have also been developed for combinatorial delivery of antibiofilm and antimicrobial agents to maximize their therapeutic effect^11–13^. Beyond their role as drug delivery vehicles, nano- and biomaterial platforms designed to modulate bacterial biofilm formation are highly desirable to enhance their therapeutic effects against chronic infections^14^.

During biofilm formation, EPS are produced to aid in bacteria surface attachment, colonization, and propagation^1,15^. Electrostatic interactions between the cationic EPS and charged antimicrobial agents prevent penetration, inhibit antibiotic activity, and confer antibiotic resistance^16^. In this way, EPS mediates biofilm resistance to charged therapeutic agents including antibiotics and antimicrobial peptides^17,18^. Shifts to investigate alternative agents, such as enzymes, for antibiofilm therapeutics are of interest^19,20^. Studies have previously reported the antibiofilm ability of the hydrolase, α-amylase, to disperse *P. aeruginosa* biofilms and inhibit biofilm growth by degrading the polysaccharides present in EPS within the biofilm matrix^21,22^. Another target of interest for antibiofilm therapies is extracellular DNA as it plays an extensive role in biofilm formation and structure. Within mature biofilms, extracellular DNA is produced by bacteria to stabilize the biofilm matrix^4,23,24^. Treatments with DNase I are capable of targeting and cleaving the extracellular DNA to increase mature biofilm dispersal and inhibit biofilm formation^24,25^.

In prior work, mucins and their associated glycans have been shown to downregulate genes associated with bacterial virulence^26,27^. Additional research has shown that mucins are capable of disrupting biofilms by dispersing surface-attached bacteria and have potential in modulating macrophage phagocytic activity to clear bacteria^27,28^. In our previous work, we developed a mucin-based synthetic mucus (SM) biomaterial for the delivery of antimicrobial peptides against *P. aeruginosa* (PAO1)^29^. In this prior work, we found SM biomaterials enhanced and prolonged the activity of LL-37 antimicrobial peptides against PAO1. Based on these studies, we sought to determine if SM biomaterials could potentiate the effectiveness of therapeutics against *P. aeruginosa* biofilms. We hypothesized a combinatory effect of the SM hydrogels loaded with antibiofilm agents whereby mucins could aid in bacterial dispersion and enzymatic treatment could degrade the biofilm matrix. Towards this end, we investigated the antibiofilm activity of SM hydrogels in combination with the antibiofilm agents, α-amylase and DNase. In addition to their direct impact on biofilm viability, we assessed the architecture of *P. aeruginosa* biofilms to further analyze the combined effects of enzymatic and SM biomaterial treatment.

## Methods

### Preparation of synthetic mucus hydrogels

Porcine gastric mucin (PGM) was obtained using a method to extract mucins from native porcine small intestinal tissues that we have described previously^29^. A 10 kDa 4-arm-PEG-thiol (Layson Bio) crosslinker was utilized to aid disulfide bond driven gelation with mucin. In a physiological buffer solution (154 mM NaCl, 3 mM CaCl_2_, and 15 mM NaH_2_PO_4_ at pH 7.4), 4% (w/v) PGM was mixed in a vial for 2 hours. Salts within the buffer aid in solubilizing the PGM within solution ^30^. In a separate vial, 8% (w/v) 4-arm PEG-thiol was prepared in physiological buffer. The PGM and PEG-thiol solutions were mixed at a 1:1 volume ratio to produce the SM hydrogel solution at a final composition of 2% (w/v) PGM and 4% (w/v) PEG-thiol. Hydrogel discs were prepared at 20 μL volumes in cylindrical molds. The hydrogel solution was incubated for 12 hours at room temperature to allow for gelation. To prepare loaded hydrogels, α-amylase (from *Aspergillus oryzae*, Sigma-Aldrich) or DNase I (from bovine pancreas, Sigma-Aldrich) was directly added into the PGM solution at initial concentrations of 300 U/mL and 300 U/mL, respectively. This solution was stirred for 2 hours at room temperature to ensure thorough mixing. The SM hydrogel solution was prepared as described previously resulting in final concentrations of 150 U/mL of α-amylase or 150 U/mL of DNase I based on the minimum inhibitory concentrations^21,24,29^.

### Microrheology of synthetic mucus hydrogels

Carboxylate-modified 100 nm fluorescent polystyrene (PS) nanoparticles (PS-COOH) were coated with 5 kDa PEG via a carboxyl-amine linkage as described in previous work.^30–32^ For multiple particle tracking microrheology studies, 25 μL of the gel was mixed with 1 μL of 0.002% w/v nanoparticles and allowed to equilibrate for 30 minutes prior to imaging. Nanoparticle movement was then imaged in real-time using a Zeiss 800 LSM microscope with a 63× water-immersion objective and Zeiss Axiocam 702 camera at a frame rate of 33.33 Hz for 10 seconds. Particle trajectories and diffusion rate were determined using MATLAB image analysis software. Time-averaged mean squared displacement (MSD) as a function of lag time was calculated from these trajectories. The generalized Stokes-Einstein relation was then used to determine the viscoelastic properties and network pore size of the hydrogels as described previously.^32,33^

### Preparation of biofilms

*Pseudomonas aeruginosa* (PAO1, ATCC 15692) was cultured in lysogeny broth (LB, Sigma-Aldrich) from frozen stock. The optical density was measured and a working concentration of 0.1 at OD_600_ was prepared. This working concentration was used for all bacteria and biofilm preparations. In 96 well plates, 100 μL of PAO1 per well was transferred. The plates were then covered with a Breathe-Easy sealing membrane (Millipore Sigma) to allow for gas exchange and incubated at 37°C for 24 hours to form mature PAO1 biofilms.

### Biofilm viability, growth, and disruption

Biofilm growth was assessed by quantifying the viability of PAO1 biofilms that formed after planktonic bacteria was treated with the SM hydrogel. In 96 well plates, 100 μL/well of PAO1 in LB was added. The prepared SM hydrogels were removed from the cylindrical molds and placed into the wells. Untreated biofilms were used as the controls. Plates were covered with Breathe-Easy sealing membranes and incubated for 24 hours at 37°C. After incubation, the wells were gently washed with phosphate buffer saline (PBS, Sigma-Aldrich) three times to remove any planktonic bacteria. Fresh PBS was added into each well and sterile wooden applicator sticks were then used to scrape the biofilm off the walls of each well. The PBS containing the scraped biofilm was collected and made into serial dilutions of 1×10^5^ to 1×10^8^ for spot plating via 10 μL spots onto agar plates. The plates were incubated at 37°C for 24 hours and the colony forming units (CFUs) were counted. Based on the plated volume, the logarithm (base 10) of CFU/mL was calculated and used to quantify PAO1 viability for biofilm growth. Biofilm disruption was measured by quantifying the viability of PAO1 biofilms after mature 24 hour old biofilms were treated with SM hydrogel conditions. Mature biofilms grown in 96 well plates and gently washed with PBS to remove planktonic bacteria. Fresh LB was added to the wells at 100 μL/well and the hydrogels were added. SM hydrogel treatment, biofilm collection, bacteria plating, and PAO1 viability were repeated and measured as described previously.

### Cotreatment and pretreatment of biofilms with antibiofilm agents

Biofilms were cotreated with free α-amylase or free DNase I in combination with SM hydrogels to assess the effect on biofilm disruption. Mature PAO1 biofilms were prepared as described prior. LB with 150 U/mL α-amylase or 150 U/mL DNase I were added to the wells at a total volume of 100 μL/well, and then SM hydrogels were added. Incubation conditions, biofilm collection, bacteria plating, and PAO1 viability were repeated and measured as described previously. Biofilm disruption was also assessed for pretreatment with free α-amylase or free DNase I prior to SM hydrogel addition. For pretreatment, mature PAO1 biofilms were first treated with 150 U/mL α-amylase or 150 U/mL DNase I in LB at a total volume of 100 μL/well and incubated for 24 hours at 37°C. The wells were washed with PBS and replaced with 100 μL of fresh LB in which the SM hydrogels were then added. The plates were incubated for an additional 24 hours at 37°C. Biofilm collection, bacteria plating, and PAO1 viability were repeated and measured similarly.

### Biofilm culture on air-liquid-interface tilted-plate

PAO1 forms biofilms on surfaces at the air-liquid-interface (ALI). Due to the nature of PAO1 biofilms to form at the ALI, an alternative method for growing biofilms on well plate bottoms for fluorescent microscopy imaging is needed. An ALI tilted-plate set-up was used to grow mature PAO1 biofilms. Using a tilted-plate set-up, the ALI can be aligned with the well plate bottom, and hence, allows the biofilm to grow at the well plate bottom for more accessible microscopy imaging. Optical black 96 well plates were tilted to a roughly 45° angle using wooden applicator sticks. After attaching the sticks to the plates, 40 μL of green fluorescent protein (GFP) expressing PAO1 (ATCC 10145GFP) culture was added into each well at a working concentration of 0.1 at OD_600_. The plates were incubated for 24 hours at 37°C to form mature biofilms on the well plate bottom. Afterwards, the plates were gently washed with PBS three times to remove any planktonic bacteria. Fresh LB was added at 40 μL/well and SM hydrogels were added. For hydrogel treatment, the plates were incubated again for 24 hours at 37°C. After incubation, the media was removed and gently washed with PBS three times. After hydrogel treatment, the biofilms were stained with 1 μg/mL of DAPI in PBS. The matrix of mature biofilms is partially composed of extracellular DNA. Thus, staining with DAPI will allow the visualization of the biofilm matrix and GFP will indicate bacteria. Plates were incubated with DAPI for 15 minutes at 37°C and then washed with PBS. A volume of 100 μL of fresh PBS was added into each well for microscopy imaging.

### Fluorescent confocal microscopy for biofilm thickness and surface area attachment

Biofilms grown on ALI tilted-plates were imaged post SM hydrogel treatment via confocal fluorescence microscope (Zeiss Confocal LSM 800) fitted with a 25x water-immersion objective. DAPI and GFP were imaged three times per well using 385 nm and 475 nm lasers, respectively. Using the GFP channel, the top and bottom Z-heights for the biofilms were determined. The difference between the Z-heights was used to calculate the optical thickness of the biofilms. Z-stack images were taken and analyzed using ImageJ analysis software to transform into sum slice Z-projection images. These projections were then converted into binary images to calculate the percent surface area of biofilm attachment.

### Statistical analysis

Data were graphed and statistically analyzed using GraphPad Prism 9. Bar graphs show means with standard deviations, dot plots show means, and box and whisker plots show medians with 5th to 95th percentiles. For analysis between multiple groups, a one-way ANOVA with Tukey’s post hoc correction was performed. Values of *P*<0.05 were considered statistically significant.

## Results

### Treatment of PAO1 biofilms with DNase and α-amylase loaded synthetic mucus biomaterials

PAO1 viability was quantified via log_10_(CFU/mL) to determine the antibiofilm activity of the SM hydrogels and SM hydrogels loaded with antibiofilm agents, α-amylase or DNase I, on biofilm growth and disruption. Particle tracking microrheology confirmed successful formulation of α-amylase and DNase I loaded SM hydrogels which remained stable up to 7 days (**Fig. S1**). The treatment of planktonic PAO1 and assessment of the viability of PAO1 in the resulting biofilm formed was used to assess the effect of SM treatment on young biofilm growth (**Fig. 1A**). A significant decrease in PAO1 biofilm viability was observed after treatment of planktonic PAO1 with SM hydrogel alone as well as DNase-loaded SM hydrogels compared to that of the untreated control (**Fig. 1B**). However for the α-amylase conditions, loading α-amylase into SM hydrogels (α-amylase-SM) did not show any significant difference compared to free α-amylase or the untreated conditions. Conversely, DNase-SM demonstrated a decrease in PAO1 biofilm viability compared to free DNase treatment only. Our data indicate that treatment with SM hydrogels and SM hydrogels loaded with DNase I can reduce initial formation and growth of young biofilms. The treatment of mature biofilms and quantification of the resulting PAO1 biofilm viability was used to measure biofilm disruption (**Fig. 1C**). SM and all α-amylase treatment conditions showed no significant differences in PAO1 biofilm viability compared to the untreated control. Treatment with free α-amylase, α-amylase-SM, and free DNase I resulted in a significant increase in PAO1 viability (**Fig. 1D**). Overall, DNase-SM showed the most significant decrease in PAO1 biofilm viability compared to α-amylase, SM, and untreated conditions.

**Figure 1.**
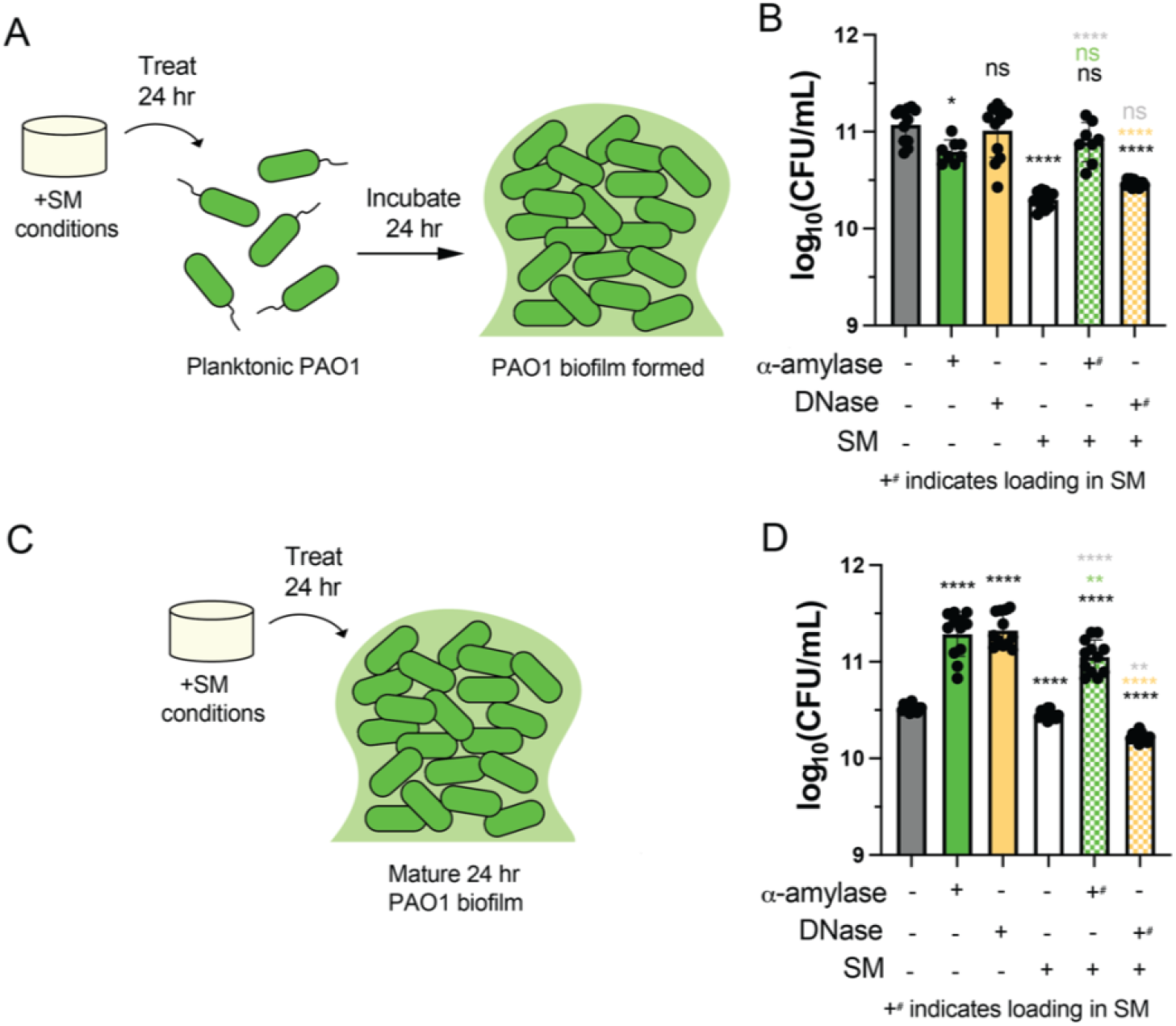
Biofilm growth and disruption after treatment with antibiofilm agents and synthetic mucus biomaterials. (**A**) Diagram of SM condition treatment on planktonic PAO1 and subsequent biofilm growth to assess biofilm growth. (**B**) PAO1 biofilm viability after 24 hour SM hydrogel treatment of planktonic PAO1 (*n* = 12). (**C**) Diagram of SM condition treatment on mature 24 hour old PAO1 biofilms to assess biofilm disruption. (**D**) PAO1 biofilm viability after 24 hour SM hydrogel treatment of mature 24 hour old PAO1 biofilms (*n* = 12). \**P*<0.05, \*\**P*<0.01, and \*\*\*\**P*<0.0001 for one-way ANOVA. Black, green, and orange asterisks indicate comparison to the untreated control, free α-amylase, and free DNase, respectively.

### Alterations to biofilm architecture following treatment with synthetic mucus biomaterials

GFP expressing PAO1 (GFP-PAO1) were used to form mature biofilms on ALI tilted plates and were imaged via confocal fluorescence microscopy (**Figure 2**). Mature PAO1 biofilms possessed a porous matrix structure and were populated by colonies of GFP-PAO1. The percent biofilm surface area was calculated from the fluorescent z-stack projection images (**Figure 2B-D**). Mean optical thickness of the biofilms was measured from orthogonal images obtained (**Figure 2E**). Using this approach, biofilm surface area and thickness were evaluated following treatment with antibiofilm agents and SM hydrogels (**Figure 3**). We observed a visual disruption in the biofilm matrix following treatment with SM hydrogels alone and in gels loaded with α-amylase and DNase (**Fig. 3B,D,F)**. Free DNase and α-amylase treatments did not appear to substantially disrupt the biofilm matrix (**Fig. 3C,E)**. Compared to untreated controls, measured percent surface area was significantly reduced for biofilms treated with SM hydrogels and α-amylase loaded SM hydrogels (**Fig. 3G)**. Biofilm thickness was significantly reduced following treatment with free α-amylase and SM hydrogels (**Fig. 3H)**. Loading of DNase and α-amylase into SM hydrogel led to further enhancements in biofilm dispersal based on their measured thickness.

**Figure 2.**
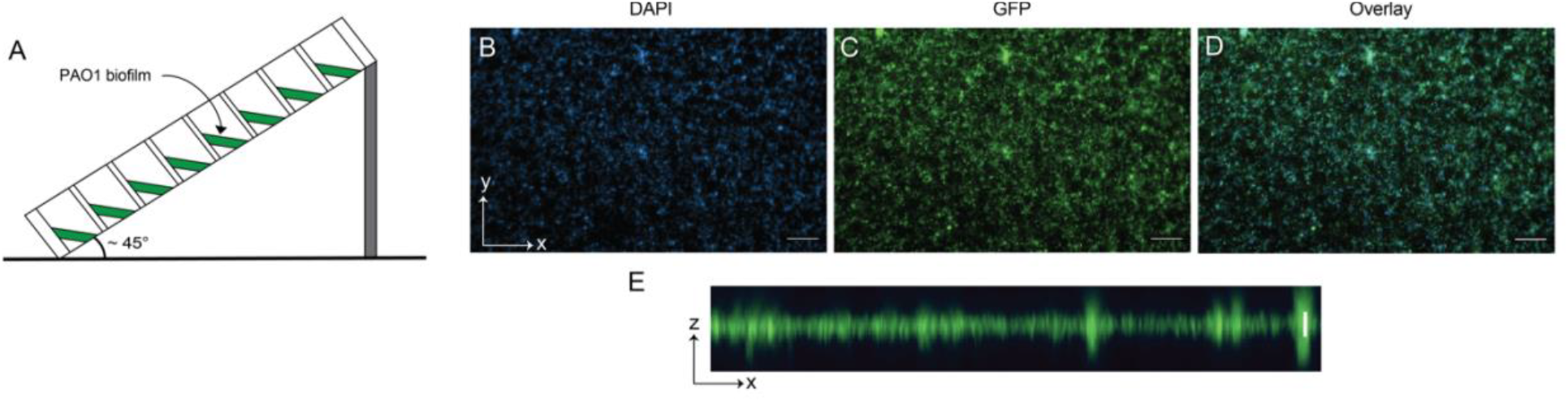
Imaging biofilm surface area attachment and thickness via confocal microscopy. (**A**) Schematic of biofilm preparation and growth on a tilted optical plate for imaging. (**B-D**) Z-stack projections of GFP-PAO1 within the biofilm matrix stained with DAPI. Scale bar = 50 μm. (**E**) Representative orthogonal cross-section of PAO1 biofilms. Scale bar = 20 μm.

**Figure 3.**
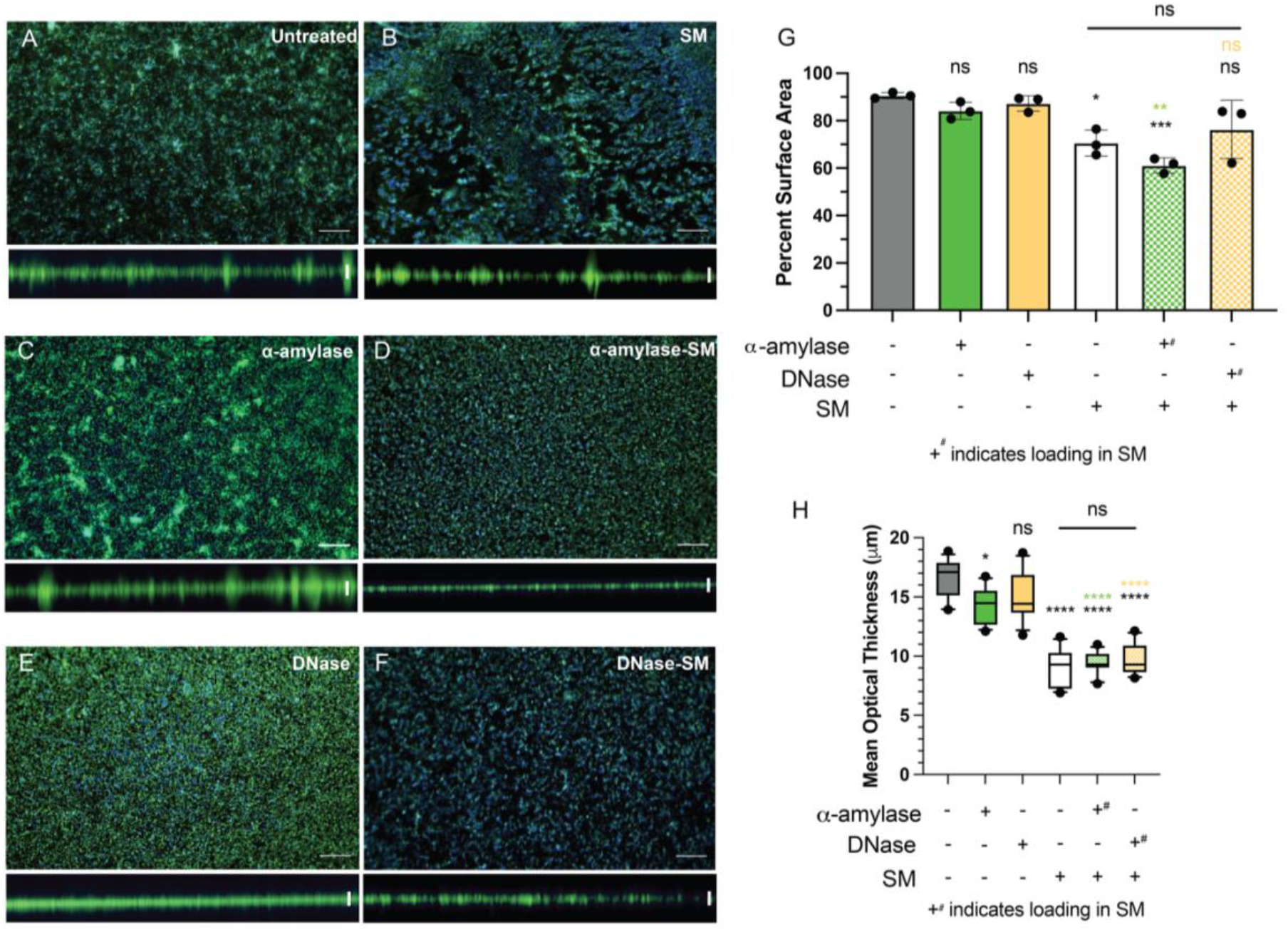
Alterations to biofilm architecture following treatment with antibiofilm agents and synthetic mucus biomaterials. (**A-F**) Following treatment with free antibiofilm agents and/or SM hydrogels, mature 24 hour old PAO1 biofilms were imaged via fluorescent confocal microscopy. GFP-PAO1 indicate viable bacteria and DAPI stained the biofilm matrix. Representative Z-stack projections (scale bar = 50 μm) and orthogonal cross-sections (scale bar = 20 μm) of the biofilm matrix are shown for each condition. (**G**) Percent surface area biofilm coverage after 24 hour treatment of antibiofilm agents and/or SM hydrogel (n = 4). (**H**) Mean optical thickness of biofilms after 24 hour treatment of antibiofilm agents and/or SM hydrogels (n = 4). \**P*<0.05, \*\**P*<0.01, \*\*\**P*<0.001, and \*\*\*\**P*<0.0001 for one-way ANOVA. Black, grey, green, and orange asterisks indicate the comparison to the untreated control, α-amylase treated biofilms, and DNase treated biofilms, respectively.

### Combined and sequential treatment with antibiofilm agents and synthetic mucus biomaterials

The effect of the timing of treatment with α-amylase, DNase I, and SM hydrogels on PAO1 biofilm viability were investigated. Cotreatment of mature biofilms included treatment with free α-amylase or free DNase I simultaneously with SM hydrogel conditions (**Figure 4A**). Compared to the untreated control, treatment with only α-amylase or only DNase did not result in any significant differences in PAO1 biofilm viability. Cotreatment with free α-amylase and SM showed significant decreases in PAO1 viability compared to the untreated control (**Figure 4B**). However, cotreatment with free DNase and SM hydrogel did not decrease CFU/mL compared to the untreated control and free DNase treatment alone. Pretreatment of biofilms consisted of treating mature biofilms with free α-amylase or free DNase I for 24 hours prior to SM hydrogel treatments (**Figure 4C)**. In comparison to the untreated control, α-amylase pretreatment followed by SM hydrogel treatments resulted in significant decreases in PAO1 viability (**Figure 4D**). In addition, pretreatment with the antibiofilm agents alone did not demonstrate any significant differences in biofilm viability. Pretreatment with DNase followed by SM hydrogel treatment did not significantly affect PAO1 biofilm viability compared to both the antibiofilm agent alone and the untreated control.

**Figure 4.**
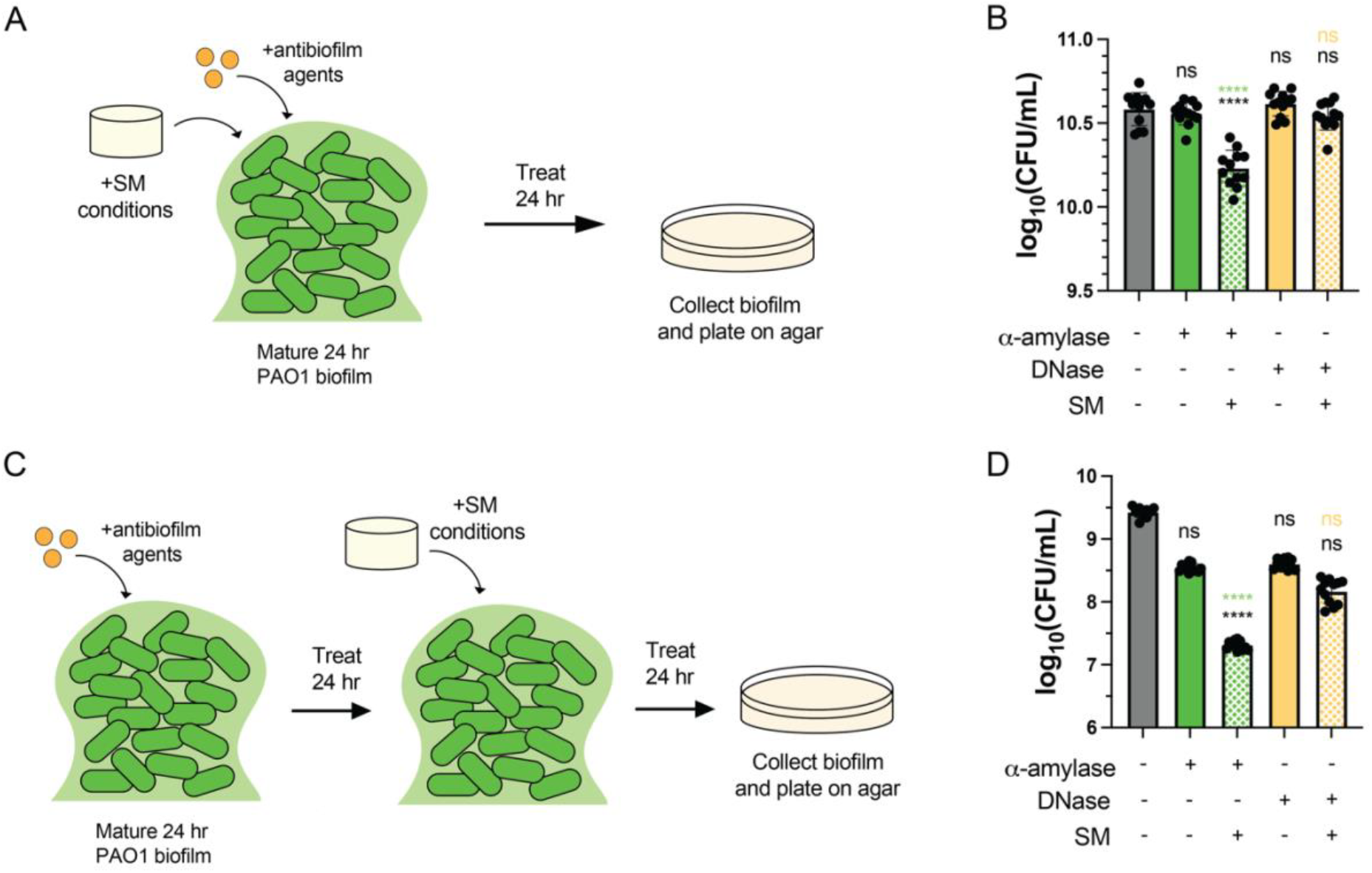
Biofilm disruption after combined and sequential treatment with antibiofilm agents and synthetic mucus biomaterials. (**A**) Schematic for the cotreatment of planktonic PAO1 with α -amylase or DNase to assess the effect on biofilm formation. (**B**) PAO1 viability after 24 hour cotreatment of with α-amylase or DNase in combination with SM hydrogels (*n* = 12). (**C**) Schematic for the pretreatment of mature 24 hour old PAO1 biofilms with α -amylase or DNase to assess the effect on biofilm disruption. (**D**) PAO1 viability after 24 hour pretreatment with α-amylase or DNase prior to 24 hour SM hydrogel treatment of mature 24 hour old PAO1 biofilms (*n* = 12). \**P*<0.05 and \*\*\*\**P*<0.0001 for one-way ANOVA. Black, green, and orange asterisks indicate the comparison to the untreated control, α-amylase treated, and DNase treated, respectively.

## Discussion

We found through these studies SM biomaterials can reduce biofilm growth and disrupt established biofilms as measured by PAO1 viability within the biofilm. Mature PAO1 biofilms range in thickness between 15 and 20 μm^34,35^. Up to ∼50% reductions in biofilm thicknesses were observed following treatment with the SM biomaterials. This is consistent with prior literature showing *P. aeruginosa* biofilms can be dispersed by mucins by triggering active biofilm bacterial escape through the detachment of cells from the biofilm matrix surface^28,36^. In addition, *P. aeruginosa* has been previously shown to adhere to native mucins and, therefore, prevent the aggregation of planktonic bacteria to limit biofilm attachment and growth^26,27^. Thus, we attribute the inhibition of biofilm growth to blocking bacterial surface attachment. Disruption of existing biofilms following treatment with SM hydrogels is likely the result of bacteria dispersal from the biofilm matrix and not due to direct bacterial killing.

The addition of the antibiofilm agents, α-amylase or DNase, to SM hydrogels also demonstrated a similar ability to disrupt biofilms. Significant reductions in biofilm surface area attachment and thickness were observed following treatment with enzyme-loaded SM hydrogels. However, this did not appear to have an additive effect with the intrinsic biofilm-disrupting properties of SM biomaterials. It should be noted when these agents are incorporated into SM biomaterials, they must be slowly released over time to disrupt the biofilm matrix. This may explain, in part, the modest reductions in biofilm viability following treatment with α-amylase or DNase-loaded SM biomaterials. Studies to account for differences in the release rate of these agents from SM hydrogels would also be helpful to determine how this may impact their performance. In addition, we found treatment with α-amylase or DNase alone can potentially increase biofilm viability. This may be the result of polysaccharides and extracellular DNA released from the biofilm matrix acting as a nutrient source for PAO1^37,38^.

We hypothesized a combined treatment approach may also enhance the anti-biofilm activity of α-amylase, DNase, and SM biomaterials. In particular, we considered a pre-conditioning step with enzymatic degradation of the biofilm matrix could enhance SM biomaterial mediated dispersal of bacteria within the biofilm. Encouragingly, we observed greater than 2 log reductions in viability in PAO1 biofilms treated initially with free α-amylase and subsequently treated SM biomaterials. Cotreatment with free α-amylase and SM biomaterials also significantly reduced PAO1 biofilm viability. Conversely, combined or sequential treatment with DNase and SM biomaterials did not have significant impact on biofilm viability. This suggests degradation of EPS aids in mucin-mediated dispersal of bacteria from the biofilm. Pretreatment of mature PAO1 biofilms with both α-amylase and DNase may lead to further enhancement in biofilm disruption using SM biomaterials and we plan to explore this in future studies.

## Conclusions

We have demonstrated that the combinatorial treatment of SM biomaterials with α-amylase or DNase can prevent growth and effectively promote disruption of static *in vitro* PAO1 biofilms. This biomaterial strategy may be particularly suited for local treatment of biofilm-associated infections. Degradation of the biofilm matrix could also improve effectiveness of antibiotics and other antimicrobial agents. We plan to explore in future studies if SM hydrogels may facilitate antimicrobial delivery to biofilms. The results of this work provide further motivation for future *in vivo* studies using these mucin-based biomaterials in animal models of chronic infection.

## Acknowledgements

This study was supported by the NIH (EB030834, HL160540, and HL160540-S1 awarded to S.Y.), and NSF GRFP (GRFP DGE1840340 awarded to S.Y.).

