## Supplemental Information for "Synthetic mucus biomaterials synergize with antibiofilm agents to combat *Pseudomonas aeruginosa* biofilms"

### Supplementary materials

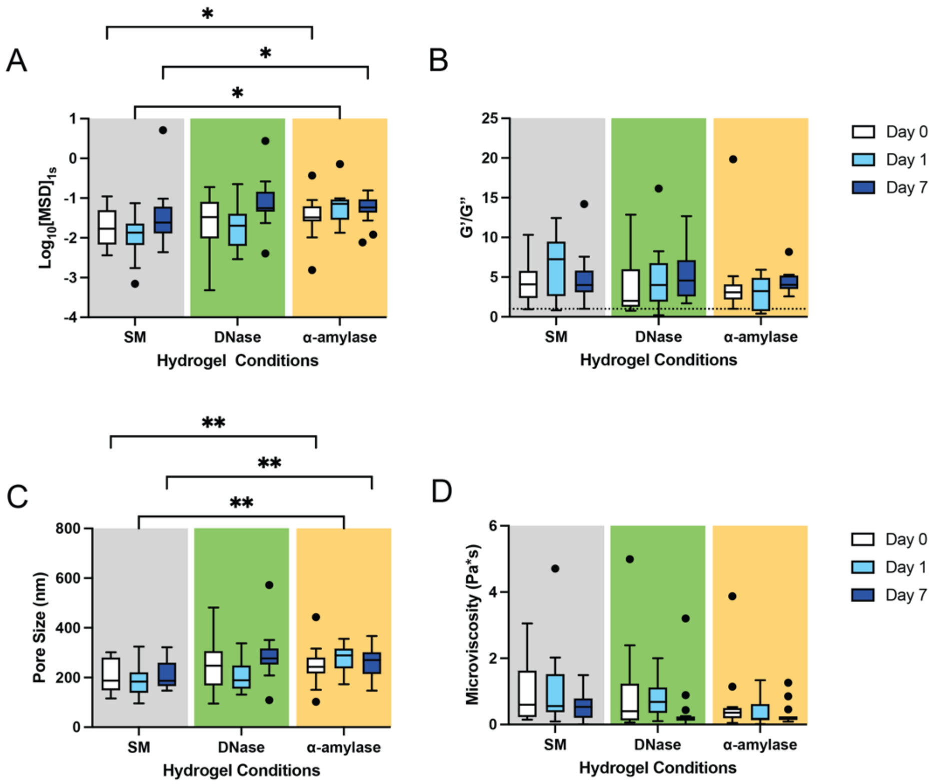


**Figure S1. Microrheological properties of SM hydrogels.** Multiple particle tracking microrheology using 100 nm particles as probes of SM, α-amylase-SM, DNase-SM hydrogels over 7 days (n = 3). (**A**) log (base 10) MSD at 1s. (**B**) G'/G" moduli ratio of SM hydrogels. (**C**) Pore size of SM hydrogels. (**D**) Microviscosity of SM hydrogels. *$P$<0.05 and **$P$<0.01 for one-way ANOVA.
